# A combinatorial approach to ALS therapies in the PrP.TDP43-315 model of ALS; complications and tribulations

**DOI:** 10.1101/2024.10.01.616092

**Authors:** Susan Fromholt, Amanda Lopez, Guilian Xu, Qing Lu, James Wymer, David R. Borchelt

## Abstract

Effective treatment for sporadic amyotrophic lateral sclerosis has been steadily advancing towards combinatorial therapies. For many years, riluzole was the only approved drug and it offered modest benefits. In 2017, edaravone was approved for ALS and became the first drug to be used in combination with riluzole. In the present study, we have attempted to build on the concepts of combinatorial therapy by testing novel drug combinations in a transgenic mouse model. Mice that express A315T mutant TDP43, using the mouse prion promoter, have been reported to develop many of the symptoms of ALS, including paralysis. Aberrant TDP43 function is a common feature in sporadic ALS, and thus TDP43 transgenic models may recapitulate disease processes that occur in humans with sporadic ALS. Although the PrP.TDP43-A315T model has been reported to develop abnormalities in gut motility that contribute to early mortality, recent studies indicated that gut motility issues could be mitigated by feeding mice with gel-based diets rather than the standard dry chow. In the present study, we have attempted to use the PrP.TDP43-A315T model to test whether we could identify a drug combination that synergized to extend life span in this model substantially. The drug combinations were built around the existing drug modalities, adding additional drugs that had indications of utility from the literature. To mitigate gut motility issues, we fed the mice gel-based diets with or without added drug combinations. Although the gel-based diets extended life expectancy in PrP.TDP43-A315T mice, most of the animals still developed gut motility abnormalities that may have contributed to early mortality. None of the drug combinations we tested extended life expectancy in this model substantially.

## Introduction

Amyotrophic lateral sclerosis (ALS) is a fatal disease of the nervous system that causes paralysis and death. The vast majority of ALS cases are classified as sporadic, meaning the cause of the disease cannot be precisely defined [1]. The sporadic form of ALS has similar pathologic features to a form of the disease caused by mutation in TARDBP-43 (TDP43) [2]. Pathologic similarities include mislocalization of TDP43 from nuclear to cytoplasmic compartments, TDP-43 immunoreactive cytoplasmic inclusions, and mis-splicing of TDP43 client mRNAs for genes such as stathmin-2 and UNC13 [3–8]. Thus, considerable evidence points towards abnormal TDP43 function as contributing to the symptoms of ALS.

For much of the past 4 decades, the prevailing approach to finding a new treatment for sporadic ALS had been to identify a candidate drug and test that one drug in ALS patients [9]. In many cases, the candidate drug was first identified to produce a modest benefit in the G93A SOD1 mouse model before testing in humans [10]. Unfortunately, this approach failed to deliver drugs that greatly extended life expectancy in humans. Here, we pursued the hypothesis that cocktails of drugs may be more effective and that such cocktails could be identified by testing in a mouse model that could be more relevant to sporadic ALS.

Working from the assumption that TDP43 dysfunction contributes to the pathogenesis of sporadic ALS, we sought to use a mouse model that could mimic TDP43 pathology of ALS. Multiple lines of TDP43 transgenic mice have been described with a range of ALS-related phenotypes. Mice that express A315T mutant TDP43, using the mouse prion promoter, have been reported to develop many of the symptoms of ALS, including paralysis [11]. The phenotype of this model is complex as abnormalities in gut motility also contribute to early mortality [12,13]. The TDP43-A315T mice were initially generated in a hybrid strain of mice before breeding to congenic in the C57BL6/J strain. Early mortality in B6.cg.TDP43-A315T mice has been described as exclusively a consequence of gut dysmotility due to loss of neurons in the myenteric plexus of the colon [14]. Herdewyn and colleagues reported that gut motility issues could be mitigated by feeding mice with gel-based diets rather than the standard dry chow [15]. In the present study, we have attempted to use the TDP43-A315T model to test whether we could identify a drug combination that synergized to substantially extend life span in this model. To mitigate the effect of strain background on phenotype, we chose to produce cohorts of C57BL6/J x FVB/NJ F1 hybrids. To further mitigate gut motility issues, we fed the mice gel-based diets. These gel-based diets also proved to be reliable approaches to deliver drug cocktails.

In formulating our drug cocktails, we used publicly available databases to identify drugs that have been through a phase 1 or phase 2 trial in ALS and had evidence in the literature of producing a modification of some aspect of ALS disease symptoms or pathology (see Table 1). We focused on drugs that should be orally available so that they could be delivered by mixing with the gel-based diets. There are 2 drugs that are currently in use in ALS, riluzole and edavarone, and there are many patients that are currently taking both of these as treatments. We used this combination as the index treatment and added additional compounds to produce novel combinatorial formulations. We identified three nutraceuticals (CoQ10, Tauroursodeoxycholic acid [TUDCA], and L-arginine) that are well-tolerated in humans and have mechanisms of action that could potentially be beneficial in ALS. TUDCA is a component of AMX0035 (Relyvrio®) that was approved for use in ALS patients before more recent data indicated a lack of efficacy. In addition to these nutraceuticals, we have identified a number of pharmaceuticals that are in use in some fashion in neurodegenerative disease, or are in some phase of testing in human ALS patients, or have been tested alone in phase 3 in ALS and found to be safe (but not particularly effective) on their own. Our hypothesis was that to achieve significant disease modification in ALS, patients will require modulation of multiple pathways simultaneously. Thus, even drugs that failed in clinical trials might be useful if combined with other drugs that modulate other pathways. Our goal with this study was not to identify combinations that produce small additive effects, but rather to identify combinations that synergize to produce profound life extensions. For many reasons, it is difficult to test combinations of drugs in humans. Although our study provides a roadmap for conducting combinatorial pre-clinical studies in mice, our findings indicate that the PrP.TDP43-A315T model is problematic as a model of ALS.

**Table 1.** Candidate drug list for combinations to be tested in TDP43 mice.

| **Table 1. Candidate drug list for combinations to be tested in TDP43 mice.** |  |  |  |
| --- | --- | --- | --- |
| **Drug Name** | **Human Dose** | **Mechanism of Action (in ALS)** | **Mouse Dose (food formulation)** |
| **Drugs currently in use for ALS** |  |  |  |
| riluzole | 50 mg/d x 2 | glutamate antagonist (reduce glutamate mediated toxicity) | 0.087 mg/g food/day |
| edaravone | 60 mg | anti-oxidant (neuroprotection) | 0.083 mg/g food/day |
| **Neuroinflammation modulators** |  |  |  |
| celecoxib | 748 mg/d | modulator of neuroinflammation | 0.67 mg/g food/day |
| ibudilast (investigational) | 100 mg/d | reduces neuroinflammation | 0.087 mg/g food/day |
| **Neuroprotectives** |  |  |  |
| TUDCA | 1g/d/ twice | anti-apoptosis (Neuroprotection) | 0.83 mg/g food/day |
| CoQ10 (Ubidecarenone) | 2.7 g/d | Mitochondrial support (neuroprotection) | 0.83 mg/g food/day |
| ropinirole (re-purpose Parkinson's disease) | 24 mg/d | dopamine agonist, neuroprotective in TDP43 iPS cell model | 0.0015 m/g food/day |
| **Metabolic modulators** |  |  |  |
| ISRIB (experimental – not currently in use in humans) | NA | Protein synthesis modulator (neuronal function) | 0.02 mg/g food/day |
| **Neuromuscular Junction/Muscle modulators** |  |  |  |
| dalfampridine (4AP) (re-purpose from multiple sclerosis) | 10 mg/d | K<sup>+</sup> channel blocker (facilitate neuromuscular junction function) | 0.0087 mg/g food/day |
| L-arginine | variable | Nitric Oxide precursor (vasodilator for muscle repair) | 0.83 mg/g food/day |
| γ-oryzanol | variable | Muscle strength | 0.083 mg/g food/day |

## Methods

### Experimental Animals

Hemizygous B6.Cg-Tg(Prnp-TARDBP*A315T)95Balo/J male mice (JAX stock# 010700) were obtained from the Jackson Laboratories and bred to female FVB/NJ mice to produce cohorts of B6/FVB F1 mice. All mice were housed on ventilated racks with automatic watering systems. Female breeders were fed ad libitum with breeder chow (2919 Teklad) with gel-based diet added for males during the interval of mating. Males were usually removed after 3 days. Offspring were weaned to cages containing standard chow (2918 Teklad) (Inotiv, West Lafayette, IN, USA). Mice were identified by PCR amplification of DNA extracted from tail biopsy following protocols provided by the Jackson Laboratories. The primer sequences used were 50 µM 9442— GGA TGA GCT GCG GGA GTT CT, 50 µM 9443—TGC CCA TCA TAC CCC AAC TG, 40 µM OIMR8744—CAA ATG TTG CTT GTC TGG TG, 40 µM OIMR8745—GTC AGT CGA GTG CAC AGT TT. At 2 – 2.5 months of age, weanlings were provided gel based diets. Two gel-based diets were compared; hazelnut-flavored MediGel^®^ Cat# 74-10-5022 (Clear H20, Inc, Westbrook, ME) and bacon-flavored NutraGel^®^ Cat# NGB-1 or NGB-2 (Bio-Serv, Flemington, NJ). All studies involving mice followed protocols approved by the University of Florida Institutional Animal Care and Use Committee.

### Pharmacological Study

In one phase of the study, we used nontransgenic C57BL6/J mice to conduct initial pharmacological studies to confirm that the drugs/nutraceuticals were orally available. In this initial phase of work, each of the drugs/nutraceuticals was delivered by oral gavage at a defined dose (Table 1). Each compound was administered after being suspended/dissolved in water containing approximately 5% carboxymethylcellulose, using a 24 to 20-gauge oral gavage. Six adult mice, 3 male and 3 female, of varied ages (6 to 18 months) were dosed with each of the drugs followed by euthanizing the animals 1 hour after dosing by CO2 asphyxiation and decapitation (blood was collected at this time). The blood was centrifuged at 800 xg for 10 min at room temperature. The plasma was separated into clean and neatly labeled tubes. Serum was analyzed by the UF CTSI Translational Drug Development Core to determine absorbance in plasma using standard mass spectrometry methods.

### Combinatorial Drug Study

We used an online Allometric scaling calculator (http://clymer.altervista.org/minor/allometry.html) to determine a dose for mice that was comparable to current usage in humans. Stock solutions (100x) of each compound to be tested were prepared. Drugs that were water-soluble were dissolved in sterile water and premixed to produce a single 100x stock. Insoluble drugs were suspended in Almond oil by vigorous vortexing and sonication. For each combination, a single mixed stock (100x) of water insoluble drugs was generated. Each cup of MediGel® held 56g of food and we added 0.56 ml of the Water and Oil drug stocks to each of the food cups, using the pipette tip to stir in the drug mixture. The consistency of MediGel® was similar to pudding and was easily mixed. In studies using NutraGel^®^, we first heated the food to 55°C to melt the gel and then mixed in the 100 X drug stocks as appropriate. The food was then allowed to cool and solidify before placing in the animal cages. A description of the drug combinations tested and the amount of drug used is provided in Table 2.

**Table 2.** Serum level of drug 1 hour after delivery by oral gavage.

| **Table 2. Serum level of drug 1 hour after delivery by oral gavage** |  |  |  |
| --- | --- | --- | --- |
| **Drug** | **Dose/mouse** | **Avg levels in serum (n = 3M,3F)** | **Range of serum levels** |
| Riluzole | 260 µg | 200 ng/ml | 86 – 240 ng/ml |
| Edaravone | 250 µg | 2824 ng/ml | 2062 – 3245 ng/ml |
| Ibudilast | 260 µg | 2.6 ng/ml | 0.7 – 8.6 ng/ml |
| Riluzole + Edaravone + Ibudilast | 2167 µg | 7904 ng/ml | 4388 – 9524 ng/ml |
|  | 2167 µg | 92551 ng/ml | 62485 – 178,557 ng/ml |
|  | 2167 µg | 1187 ng/ml | 464 – 1652 ng/ml |
| Dalfampridine | 26 µg | 72.8 ng/ml | 22 – 142 ng/ml (Females uniformly higher) |
| Ropinirole | 4.5 µg | 1.1 ng/ml | 0.5 – 1.8 ng/ml |
| Celecoxib | 2000 µg | 2864 ng/ml | 1486 – 3613 ng/ml |
| TUDCA | 2500 µg | 69.8 ng/ml | 13 – 127 ng/ml |
| CoQ10 | 2500 µg | Not detected | Not detected |

The drug-containing diets were started when mice reached 100 days and continued until mice reached humane endpoints, except Combos 5 and 8 which were discontinued after 6 months of treatment. We used a scoring system to define a humane endpoint based on the following criteria: >= 15% weight loss from baseline weight; impaired mobility limiting access to reach food or water; labored breathing, respiratory distress or cyanosis (blue tinged color); tremors, convulsions or seizures lasting more than 1 minute or occurring more than once a day; dehydration lasting over 24 hours that is unresponsive to treatment; moribund - unable to right itself. Any animal that unexpectedly developed a spontaneous tumor was euthanized. Nontransgenic littermates of treated or untreated mice were euthanized and tissues were harvested when the last transgenic animal of any given cage reached endpoint criteria.

### Immunostaining

Immunohistochemistry was performed on 5 μm sagittal paraffin-embedded brain sections according to the standard method. Briefly, tissue sections were deparaffinized in xylene and rehydrated through a graded ethanol series. Antigen retrieval was conducted using 10 mM citrate buffer (pH 6.0) in a steamer. Endogenous peroxidase activity was quenched with 0.03% H_2_O_2_. Subsequently, sections were blocked with 3% normal serum and incubated overnight at room temperature with primary antibodies. In this study, the primary antibodies included ChAT (AB144P, Millipore Sigma, goat polyclonal, 1:500), GFAP (Dako, rabbit 1:2000), Ubiquitin (Ubi-1, Encor, mouse monoclonal, 1:1000), and TDP-43 (MCA-3H8, Encor, mouse monoclonal, 1:1000). Following washing steps, biotinylated secondary antibodies (Vector Laboratories, CA, USA) were applied, succeeded by avidin-biotin complex (ABC) (Vector Laboratories, CA, USA) treatment. 3,3’-Diaminobenzidine (DAB) kit (Seracare, MA, USA) was utilized as the chromogen. Sections were then counterstained with CAT hematoxylin (Biocare Medical, CA, USA), dehydrated, and coverslipped. All incubations were carried out in a humidified chamber to prevent section desiccation and all washes used PBS with 0.05% Tween-20. Images were acquired using an Aperio digital pathology system (Leica Biosystems).

### Statistical Analysis

Statistical analysis of differences in survival between treatment groups was done using GraphPad Prism 9.5.1 and the Log-rank (Mantel-Cox) test. Analysis of motor neuron numbers in spinal cord was done using GraphPad Prism by one-way ANOVA.

## Results

We identified a combination of nutraceuticals and pharmaceuticals that are in use in some fashion in neurodegenerative disease, or are in some phase of testing in human ALS patients (Table 1). Our plan was to use different combinations of drugs to modulate multiple cellular processes simultaneously in an attempt to achieve robust disease modifying effects that slow progression in the TDP43-A315T model. To assess the oral bioavailability of our test drugs, we treated nontransgenic B6 mice by oral gavage and collected serum at 1hour for analysis. All of the test drugs were detectable in serum except CoQ10 (Table 2).

To produce cocktails of drugs with different mechanisms of action, we devised 5 combinatorial mixtures that mixed different types of drug activities (Table 3). To mitigate gut problems in TDP43-A315T mice [15], we placed mice on gel-based diets at ∼2 months of age, initially using Hazelnut MediGel^®^ diet. We expected these mice to develop ALS-like symptoms by 6 months of age [15]. In mid-study, we began to be concerned that the Hazelnut diet was not nutritionally complete enough for long-term studies and switched to a bacon-flavored NutraGel^®^ diet.

**Table 3.** Drug Combinations.

| Table 3. Drug Combinations |  |
| --- | --- |
| Combination code | Constituents |
| Combo 1 | Riluzole, Edaravone, Ibudilast |
| Combo 2 | Riluzole, Edaravone, Ibudilast, CoQ10, Oryzanol, TUDCA, L-Arginine |
| Combo 5 | Riluzole, Edaravone, CoQ10, TUDCA, L-Arginine, 4AP dalfampridine, Celecoxib |
| Combo 8 | Riluzole, Edaravone, CoQ10, TUDCA, L-Arginine, Ropinrole, Celecoxib |
| Combo 13 | Riluzole, Edaravone, Minocycline, CoQ10, Oryzanol, TUDCA, L-Arginine, Ropinrole, ISRIB, Celecoxib |

For both of the diets, most of the mice developed symptoms resembling those of the mutant SOD1 model including hindlimb weakness with paresis (Tables 4 and 5). Many also exhibited hunched posture and weight-loss, and most of these types of mice were found to have an accumulation of stool in the colon at autopsy. In the cohorts fed the MediGel^®^ diet, about half of the mice had clear evidence of extreme compaction (see Table 4). In mice with milder GI issues the dark coloration of the MediGel® diet made it difficult to assess the severity of GI abnormalities. In the cohorts fed NutraGel^®^ diet, we also observed that most of the mice had moderate to severe GI abnormalities (see Table 5). There were three TDP43 mice that had completely normal looking intestines at the time of harvest. These three mice were among the most long-lived animals of the cohort (Table 5, yellow highlight; Fig 1A). Mice fed NutraGel^®^ diets lived longer than mice fed MediGel® diets (Fig. 1B, p<0.0001). In the MediGel^®^ cohort it was clear that female mice tended to live longer (Fig. 1C, p=0.0014). The cohort of mice fed NutraGel^®^ contained too few female mice to draw firm conclusions, but here again female mice tended to live longer (Fig 1A, open symbols; Table 5). These data demonstrated that PrP.TDP43-A315T mice can live up to 14 months when fed gel diets.

**Table 4.**
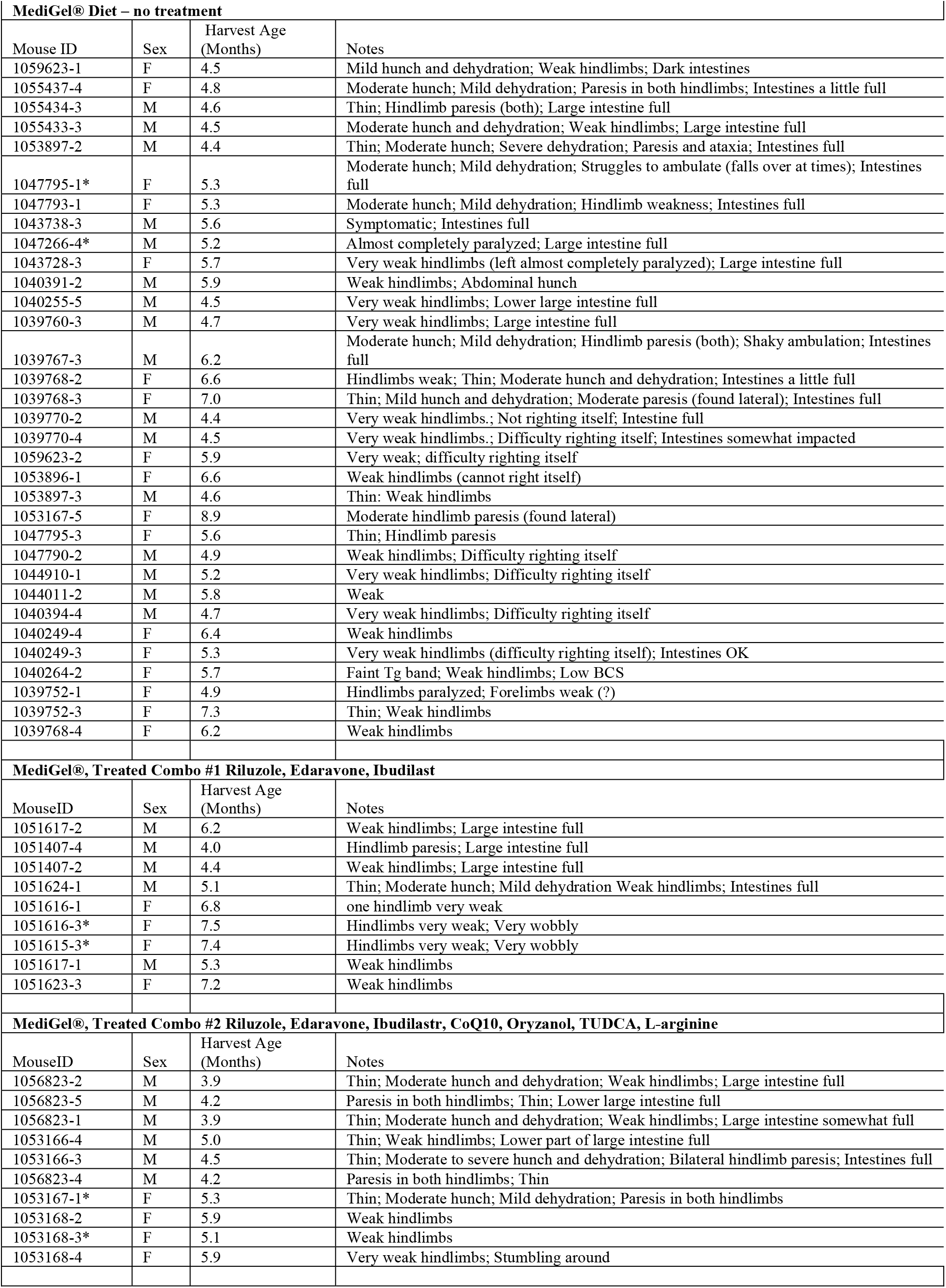
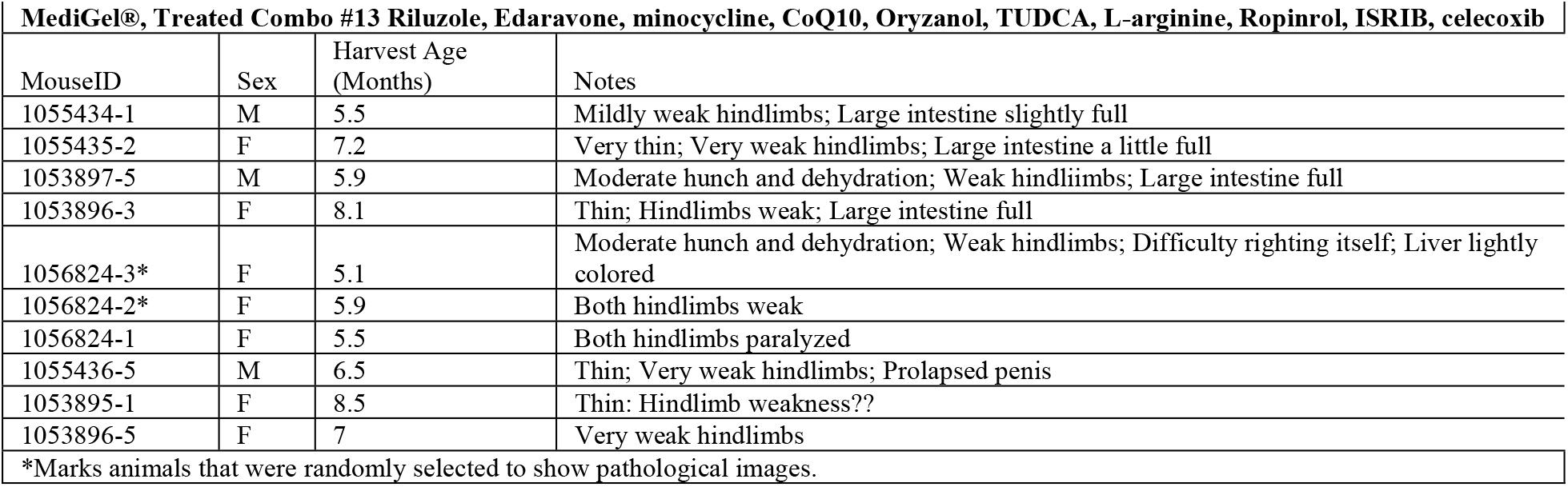
Characteristics of TDP43 mice fed MediGel® with and without drug treatment.

**Table 5.** Characteristics of TDP43 mice fed NutraGel® with and without drug treatment NutraGel® diet – no treatment.

| **Table 5. Characteristics of TDP43 mice fed NutraGel® with and without drug treatment** |  |  |  |
| --- | --- | --- | --- |
| **NutraGel® diet – no treatment** |  |  |  |
| Mouse ID | Sex | Harvest Age (Months) | Notes |
| 1111134-4 | M | 7.0 | Weak; Hunch; Thin; Intestine abnormal |
| 1111133-1 | M | 7.2 | Hindlimbs weak?; Intestines full |
| 1111165-5* | M | 5.3 | Thin; Dehydrated; Very weak hindlimbs; Intestines full |
| 1111165-4 | M | 4.7 | Thin; Weak hindlimbs and hindlimb clasp 4/29/21; Intestines very full |
| 1108345-4 | M | 7.3 | Paralyzed? (almost die); Intestines very dark |
| 1108133-1 | F | 8.5 | Very weak hindlimbs; Large intestine somewhat full |
| 1077044-5 | M | 10.8 | Moderate hunch and dehydr.; Abnormal abdom. exam; Paresis both hindlimbs (?); Intestine abnormal |
| 1077044-3 | M | 9.6 | Weak hindlimbs; Intestines full |
| 1077044-1 | M | 11.8 | Weak hindlimbs; Intestine abnormal |
| 1077073-3 | M | 11.6 | Hindlimbs very weak; Intestine abnormal |
| 1111169-2 | M | 6.0 | Hindlimb weakness?; Intestines very full |
| 1111169-1 | M | 7.2 | Hindlimbs weak; Intestines slightly full |
| 1108132-4 | M | 6.3 | Hindlimbs appeared weak; Intestines full |
| 1077044-2 | M | 8.7 | Thin; Hindlimb clasp; Paresis Intestine abnormal |
| 1077755-4* | F | 14.5 | Hindlimbs weak; Thin; Intestine clear |
| 1077043-5* | F | 12.1 | Severe paresis (difficulty righting itself); A little bit of feces but mostly clear |
| 1077043-4 | F | 11.3 | Weak; Thin; Lt. hindlimb almost paralyzed; Intestine clear |
| **NutraGel®, Treated Combo #5, off treatment after 6 months (Riluzole, Edaravone, TUDCA, CoQ10, L-arginine, celecoxib, dalfampridine)** |  |  |  |
| Mouse ID | Sex | Harvest Age (Months) | Notes |
| 1110757-4 | M | 7.7 | Weak hindlimbs?; Intestines full; |
| 1087896-1* | F | 10.6 | Weak hindlimbs; Intestines somewhat full |
| 1087901-1* | F | 11.0 | Weak hindlimbs?; Hunched; Severe dehydration; Constipated |
| 1087900-3 | M | 9.0 | Hindlimb weakness; Intestines full |
| 1087895-2 | M | 10.6 | Appeared to have abdominal hunch BUT intestines clear at harvest |
| **NutraGel®, Treated Combo #8, off treatment after 6 months (Riluzole, Edaravone, TUDCA, CoQ10, L-arginine, Celecoxib, roprinirole)** |  |  |  |
| Mouse ID | Sex | Harvest Age (Months) | Notes |
| 1110762-4 | M | 6.4 | Hindlimbs weak?; Intestines somewhat full |
| 1092415-5 | F | 10.2 | Paresis; Large intestine full |
| 1085150-5 | M | 10.2 | Weak; Hindlimb still usable; Hunch; Intestines full |
| 1085150-4 | M | 6.3 | Some hindlimb weakness; Large intestine somewhat full |
| 1110762-5 | M | 6 | Weak hindlimbs Intestine clear |
| 1085128-4* | F | 11.7 | Hindlimb paresis Intestine mostly clear |
| 1085128-2* | F | 10.3 | Hindlimb almost paralyzed Intestine mostly clear |
| *Marks animals that were randomly selected to show pathological images. |  |  |  |

**Figure 1.**
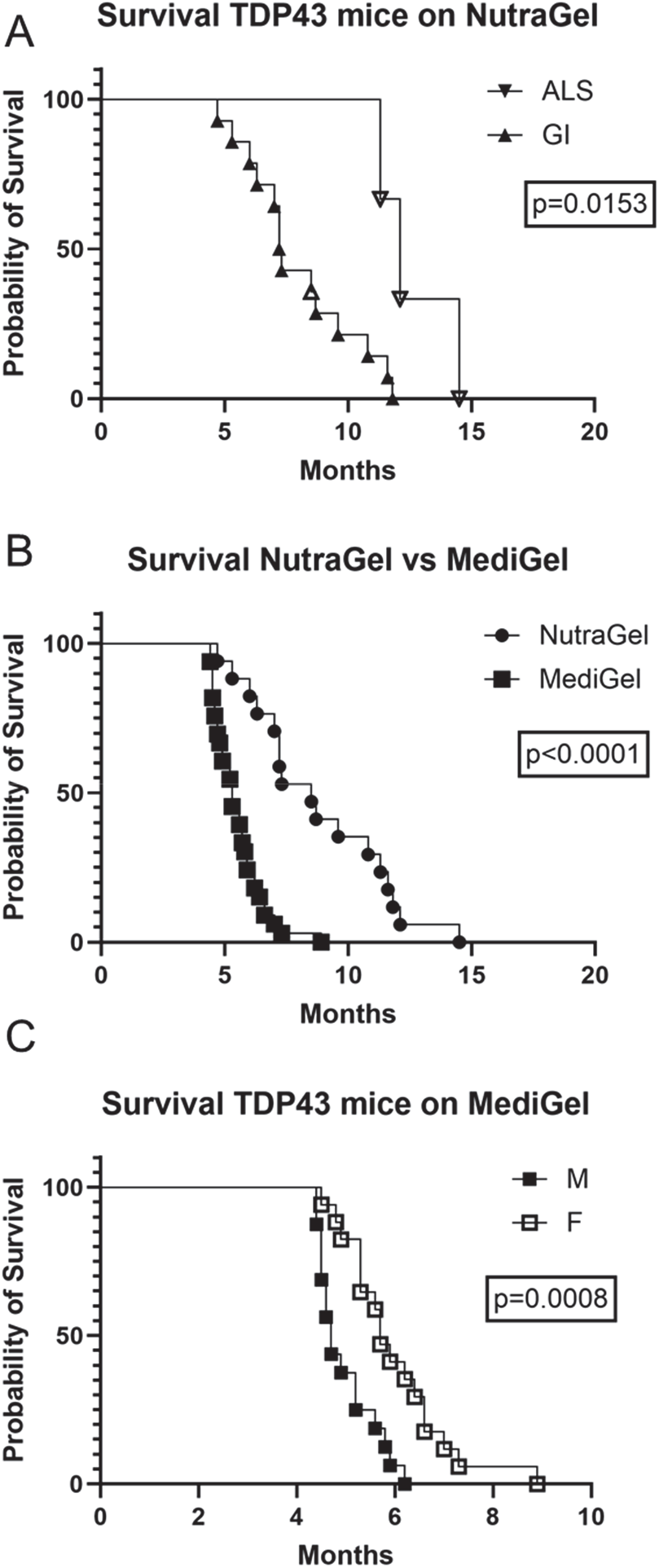
Comparison of life expectancy for TDP43 mice fed NutraGel® and MediGel®. A) The ages that TDP43 mice that were fed untreated NutraGel® reached endpoint were plotted, stratifying the data according to whether the mice exhibited GI abnormalities or only ALS-like clinical signs. The number of mice in each group were as follows: ALS-M (n = 0), ALS-F (n = 3), GI-M (n = 13), GI-F (n = 1). B) A survival plot for TDP43 mice fed NutraGel® (n = 17) versus MediGel® (n = 33) is shown. C) The ages that TDP43 mice that were fed untreated MediGel® reached endpoint were plotted, graphing males and females separately. The number of mice in each group were as follows: M (n = 16), F (n = 17). In (A) and (C), filled symbols note male mice and open symbols note females; all mice were transgenic for mutant TDP43. Some of the data points overlap and are not discernable.

To assess TDP43 pathology in these mice, we stained spinal sections with an anti-TDP43 monoclonal antibody designated 3H8. In both the transgenic and nontransgenic animals, the vast majority of TDP43 immunoreactivity was nuclear (Fig. 2). In all animals examined, there was some level of diffuse cytoplasmic immunoreactivity that was relatively variable. There were no obvious cytoplasmic inclusions in any of the animals examined or any evidence of significant loss of nuclear staining. Spinal TDP43 immunoreactivity levels, and location, in mice with obvious GI abnormalities (Fig. 2B) was not obviously different from mice that lacked obvious GI abnormalities (Fig. 2D). Overall, although there was some intensification of immunoreactivity in the TDP43 transgenic mice, there were not obvious cytoplasmic inclusions or obvious loss of TDP43 nuclear localization.

**Figure 2.**
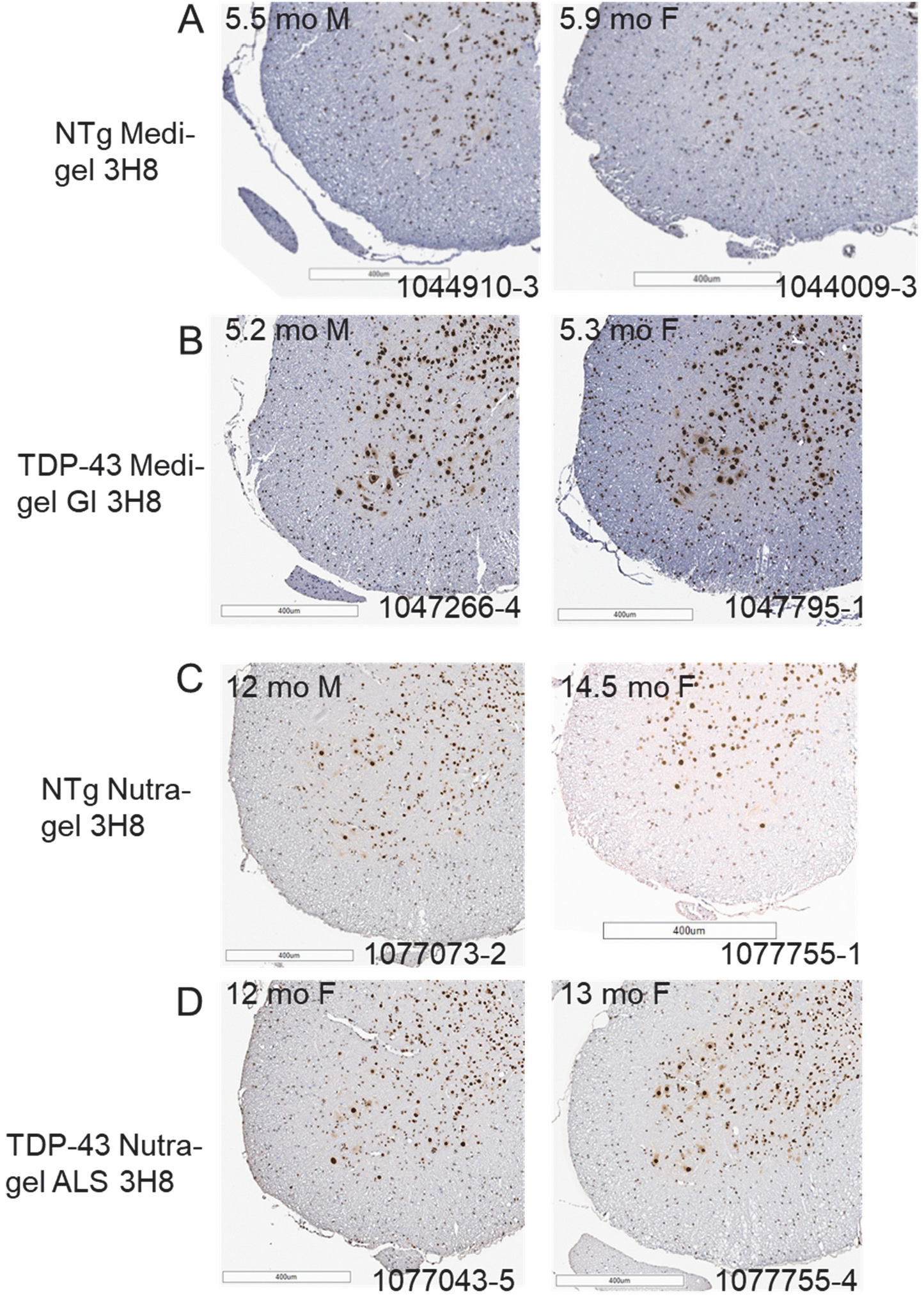
TDP43 immunoreactivity in primarily localized to the nucleus in end-stage TDP43 mice. A-D) Representative images of TDP43 immunostaining are shown. Every transgenic animal in the plots described in Figure 1, and Tables 4 and 5, was examined neuropathogically. A total of 19 nontransgenic littermates were also examined (7 from MediGel® cohort, 11 from NutraGel® cohort). The images shown reflect the most common appearance. Images of both male and female mice are shown.

To confirm that mice presenting with clinical signs resembling paralysis did indeed have motor neuron disease, we counted motor neurons in the lumbar sections of TDP43 mice fed NutraGel^®^, comparing TDP43 mice exhibiting ALS-GI and ALS symptoms with nontransgenic controls (Fig. 3). We stained sections of lumbar cord with antibodies to choline acetyl transferase (ChAT) (Fig. 3A-C) and determined the total number of positive cells in two sections from each mouse counted. The two values for each mouse were averaged and graphed (Fig. 3D). The original description of the TDP43 model reported ∼20% reductions in motor neuron numbers per section in mice that survived an average of 153 days [16]. We expected that ameliorating GI symptoms and extending lifespan might allow for more significant losses of motor neurons; however, our analysis of motor neuron numbers failed to detect significant reductions in lumbar sections (Fig. 3D). This finding indicated that any ALS-like symptoms in this model in the background strain we used were due to motor neuron dysfunction rather than motor neuron loss.

**Figure 3.**
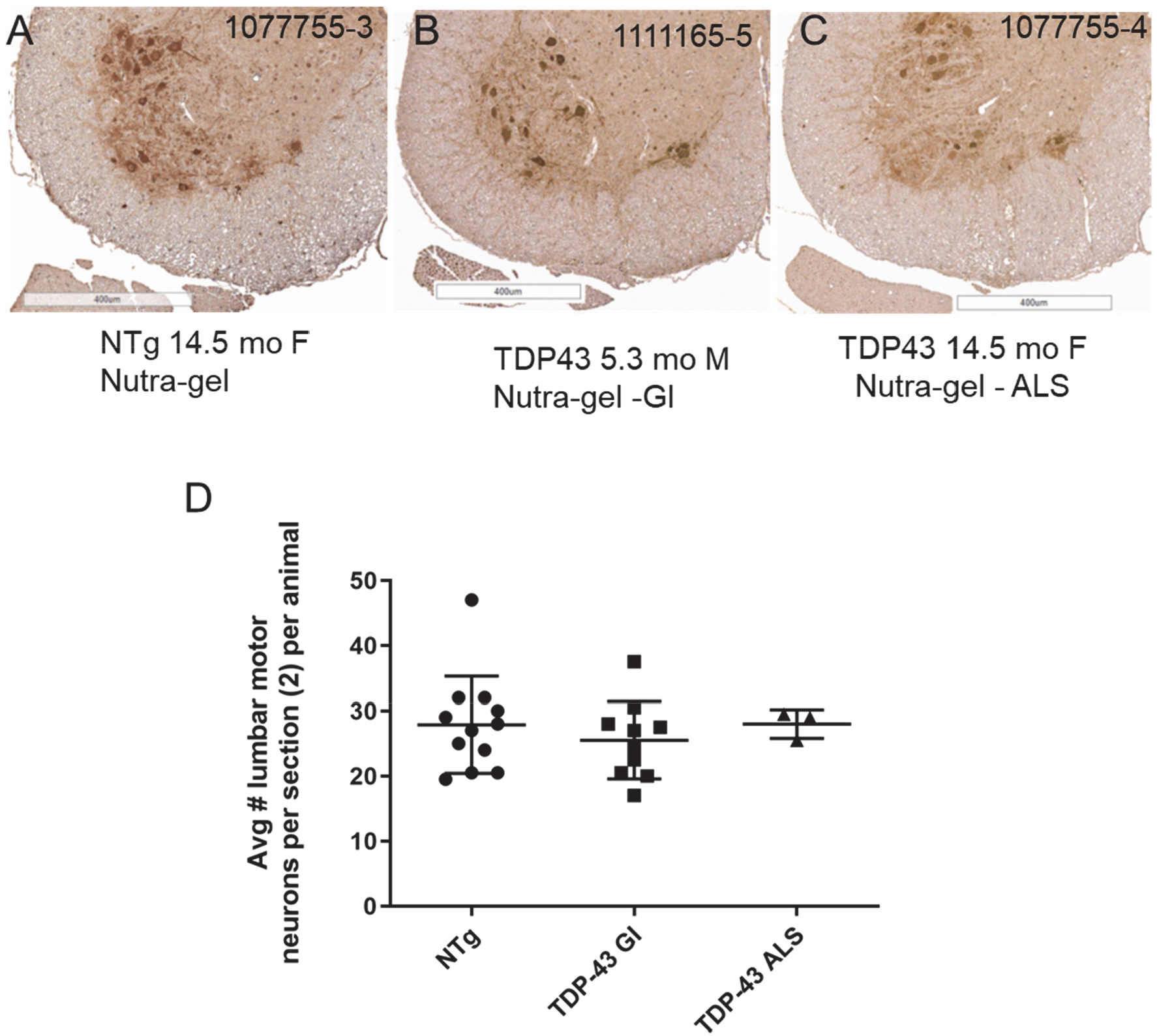
Motor neuron loss is not a prominent feature of TDP43-A315T mice. A-C) Representative images are shown of choline acetyl transferase (ChAT) immunostaining from one hemisphere of lumbar spinal cord in nontransgenic and TDP43 mice fed NutraGel®. D) For each animal examined, the number of large ChAT positive cells were counted in two sections of spinal cord. The values were averaged and plotted. The values in nontransgenic mice (n = 12; 8M; 4F) were compared to TDP43 mice, separating mice with GI abnormalities (9 M; 1F) from mice that only exhibited ALS-like clinical signs (3 F).

Regardless of whether we used the MediGel^®^ or NutraGel^®^ diets, none of the tested drug combinations showed any evidence of delaying the age to endpoint criteria substantially. In the MediGel^®^ cohort treated with Combo 1 there was no significant effect of life expectancy (Fig. 4A). TDP43 mice treated with Combo 2 tended to reach endpoint earlier than untreated mice (Fig. 4B). The only potentially promising combination was Combo 13, which appeared to modestly extend survival (Fig. 4C). In the NutraGel^®^ cohorts treated with Combos 5 and 8, neither drug combination significantly delayed endpoint criteria (Fig. 5A and B). The tendency for female TDP43 mice to live longer before reaching endpoint was also evident in these three cohorts of treated mice (Figs. 4 and 5, filled symbols – M; open symbols – F). Overall, we were not able to achieve a robust synergy for any of the combinations tested.

**Figure 4.**
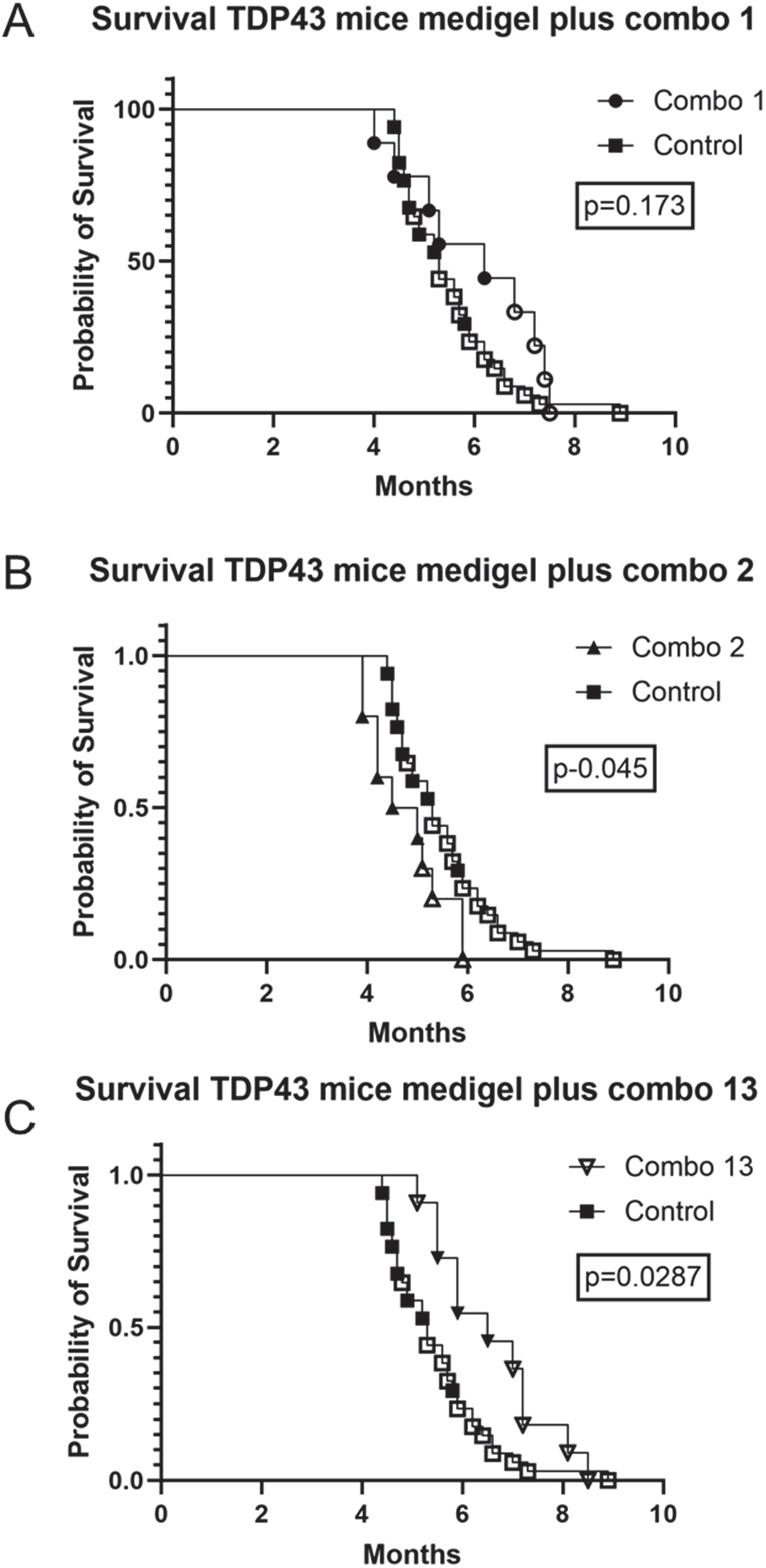
Drug combinations tested in TDP43 mice fed MediGel® failed to substantially extend survival. A) Survival plot for mice treated with Combo 1. The total number of TDP43 mice treated was nine (5 M and 4 F). B) Survival plot for mice treated with Combo 2. The total number of TDP43 mice treated was ten (6 M and 4 F). C) Survival plot for mice treated with Combo 13. The total number of TDP43 mice treated was ten (3 M and 7 F). The survival data for the treated mice was compared to a single cohort of TDP43 mice fed only MediGel® (16 M; 17 F).

**Figure 5.**
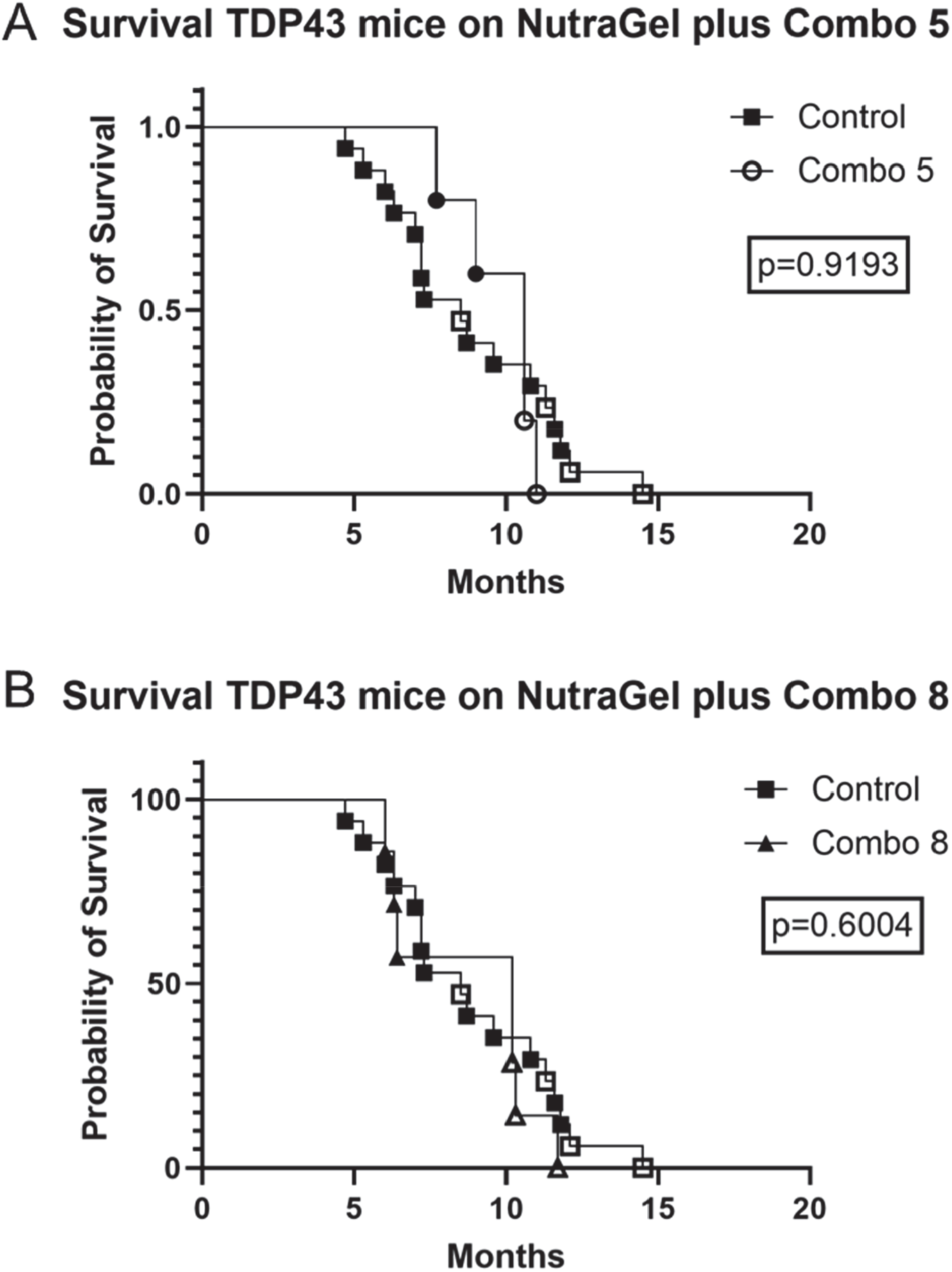
Drug combinations tested in TDP43 mice fed NutraGel® failed to substantially extend survival. A) Survival plot for mice treated with Combo 5. The total number of TDP43 mice treated was five (3 M and 2 F). B) Survival plot for mice treated with Combo 8. The total number of TDP43 mice treated was seven (4 M and 3 F). The survival data for the treated mice was compared to a single cohort of TDP43 mice fed only NutraGel® (13 M; 4 F).

In the absence of robust phenotypic improvements in survival, we asked whether any of the drug combinations may modulate pathologic abnormalities present in these mice. We assessed the levels of ubiquitin and GFAP. In most of the TDP43 mice that reached humane endpoint, we observed increased ubiquitin immunoreactivity (Fig. 6). The reactivity consisted of intensely labeled cells of which some were of a size and location to be probable motor neurons. These same types of immunoreactive profiles were observed in all of the treated mice with no dramatic change in frequency or morphology (Fig. 7). GFAP immunoreactivity in the TDP43 mice was largely indistinguishable from nontransgenic mice (Fig. 8), and there was no obvious change in the level of GFAP immunoreactivity in the spinal cords of any of the drug-treated mice (not shown). Collectively, there was no obvious indication that the drug treatments modified spinal pathology in any way.

**Figure 6.**
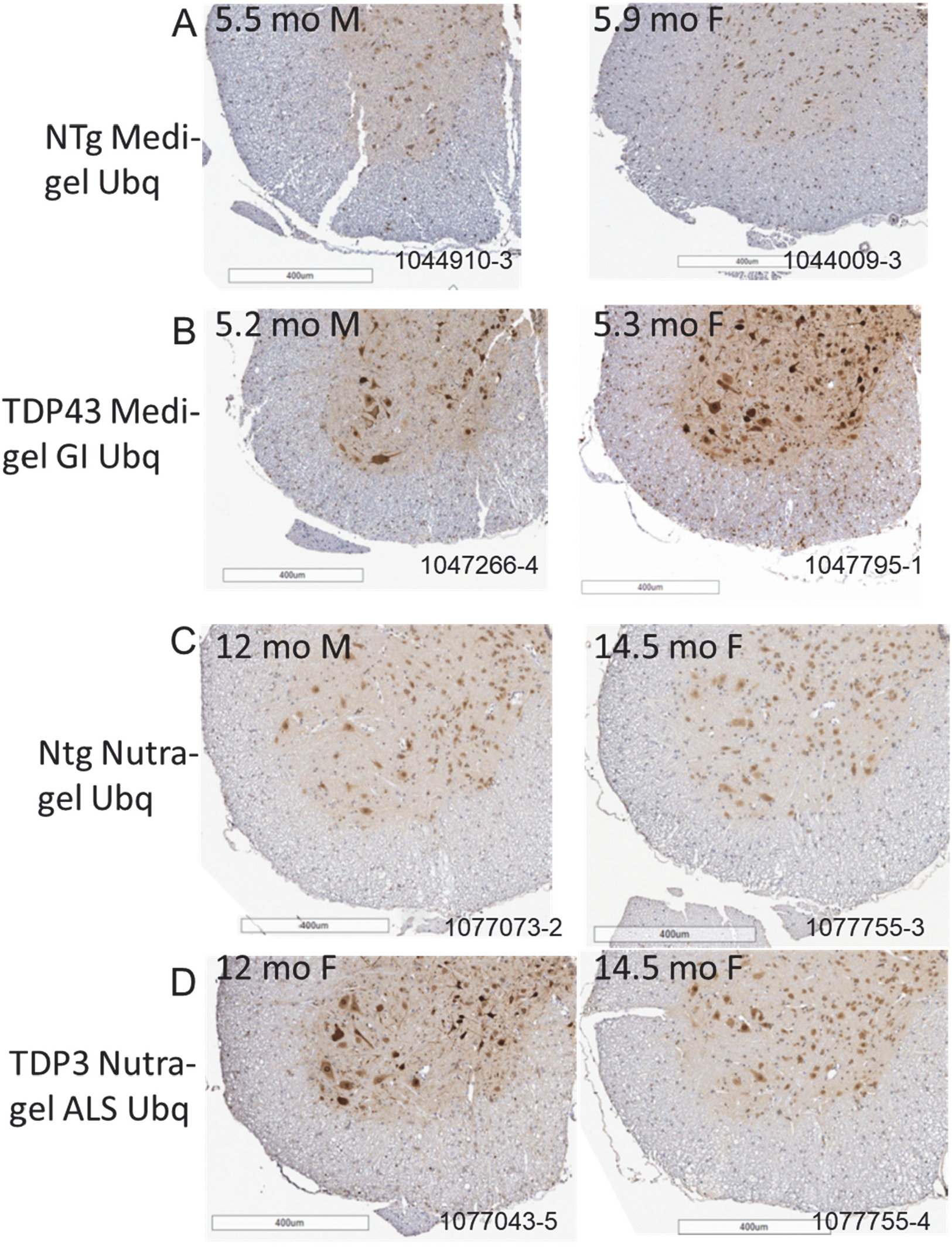
Symptomatic TDP43 mice show elevated ubiquitin immunoreactivity in lumbar spinal cord. A-D) Representative images of ubiquitin immunostaining are shown. Every transgenic animal in the plots described in Figure 1, and Tables 4 and 5, was examined neuropathogically. A total of 17 nontransgenic littermates were also examined (6 from MediGel® cohort, 11 from NutraGel® cohort). The images shown reflect the most common appearance. Images of both male and female mice are shown.

**Figure 7.**
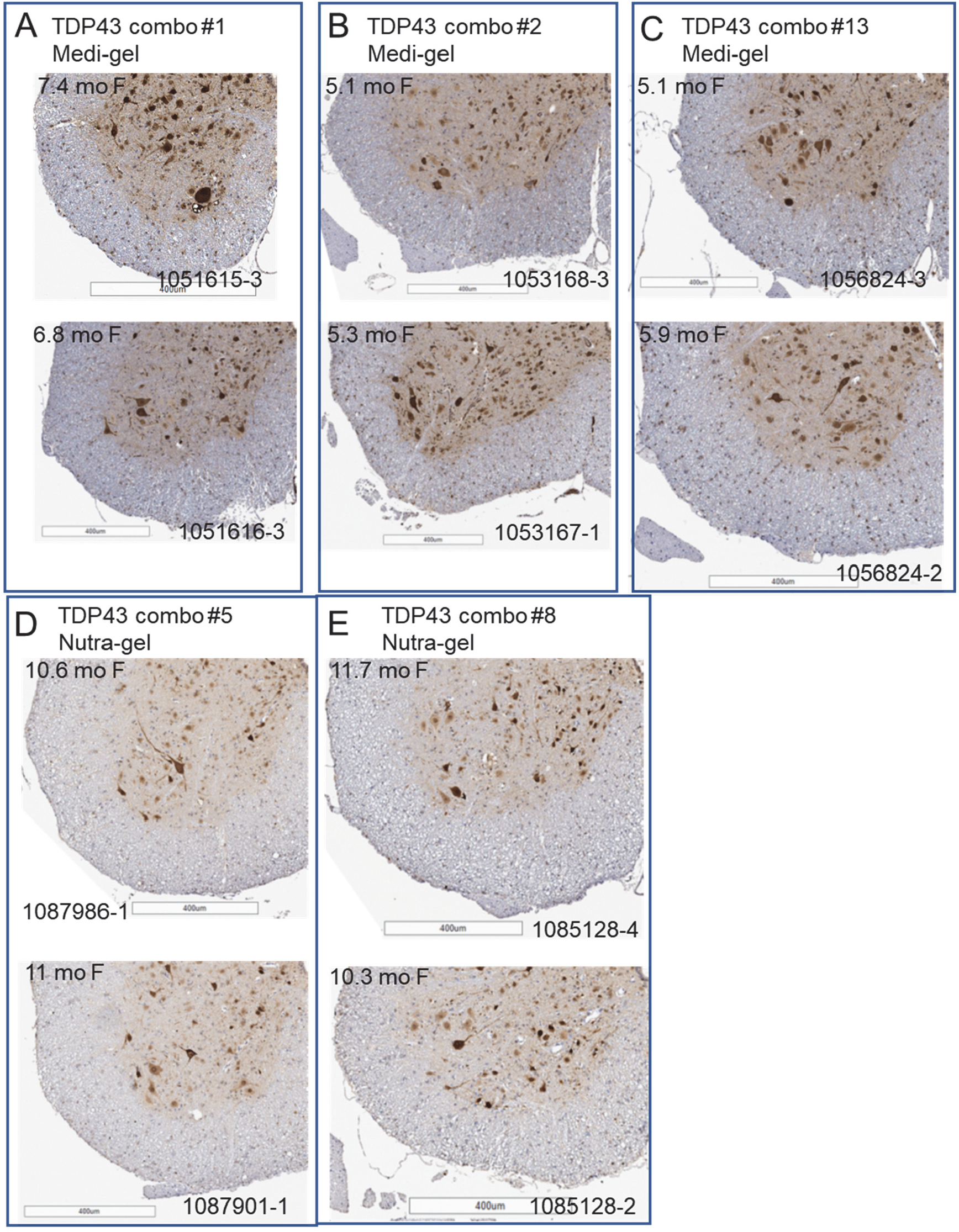
Ubiquitin immunoreactivity remains elevated in TDP43 mice treated with the drug combinations. A-D) Representative images of ubiquitin immunostaining are shown. Every transgenic animal in the plots described in Figures 4 and 5, and Tables 4 and 5, was examined neuropathogically. The images shown reflect the most common appearance. Images of both male and female mice are shown.

**Figure 8.**
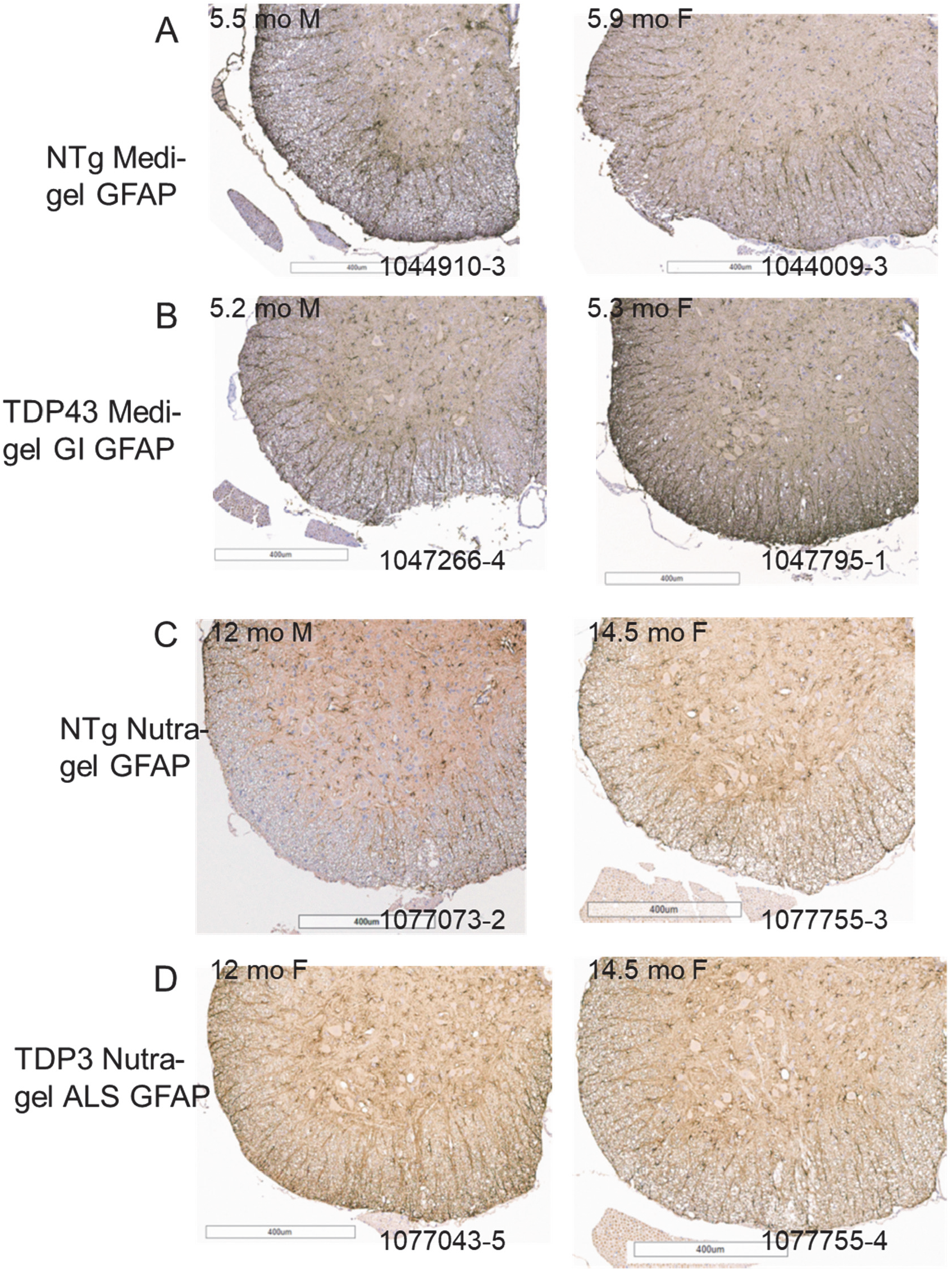
GFAP immunoreactivity in lumbar spinal cords of symptomatic TDP43 mice is largely indistinguishable from nontransgenic mice. A-D) Representative images of GFAP immunostaining are shown. Every transgenic animal in the plots described in Figures 4 and 5, and Tables 4 and 5, was examined neuropathogically. A total of 19 nontransgenic littermates were also examined (8 from MediGel® cohort, 11 from NutraGel® cohort). The images shown reflect the most common appearance. Images of both male and female mice are shown.

## Discussion

The goal of this study was to determine if we could find a combination of drugs that could act synergistically to substantially extend the survival of TDP43-A315T mice. The combinations of pharmaceuticals and nutraceuticals we tested included drugs approved for other indications that could be readily available to ALS patients. The design was to build on a foundation of Riluzole and Edaravone, which were the only 2 approved drugs at the time the study was initiated, adding candidate drugs and nutraceuticals to produce cocktails. The study included one investigational drug, Ibudilast, and one experimental drug, ISRIB, as more exploratory endeavors. Although prior studies had described GI abnormalities in the TDP43-A315T model we used [14], other studies had suggested that we could mitigate the GI symptoms with gel-based diets to allow the mice to develop ALS-like symptoms [15]. Although most of the mice did show clinical signs expected of an ALS mouse model, including weakness and hind-limb paresis, in the background stains we used here (B6/FVB F1), the gel diets did not fully mitigate GI abnormalities. We estimate that GI dysmotility directly contributed to the poor condition of most of the animals, triggering euthanasia. One of the combinations tested produced a modest delay in reaching the age to endpoint, but 4 other combinations where ineffective if not deleterious. Given the frequency of GI abnormalities in this model, it is not clear whether the TDP43-A315T model is a workable model for ALS drug discovery.

In the original description of the TDP43-A315T model, mice that were fed standard pellet chow reached humane endpoints at ages ranging from 100 to 240 days [11]. In the congenic B6 background, male TDP43-A315T mice fed normal chow reached endpoint at an average of 105 days, with females living somewhat longer to an average of 185 days [14]. When we first imported the B6.TDP43-A315T mice from Jackson Laboratories, we quickly encountered the emergence of GI abnormalities in mice fed normal pellet chow. Several of the imported breeder males succumbed to GI dysmotility before 4 months of age. Placing mice on gel-based diets at 2-2.5 months of age significantly delayed the GI symptoms and facilitated successful breeding. We quickly determined that we could easily mix cocktails of drugs into the gel diet and we identified Hazelnut flavored MediGel^®^ as a means to prolong survival and deliver our drug cocktails. We then launched an effort to test multiple combinations at the same time. As the first phase of studies progressed, we realized that the MediGel^®^ was not suitable for long-term use and identified bacon-flavored NutraGel^®^ as a more nutritionally complete gel diet. Importantly, NutraGel^®^ could be liquified at 55°C, enabling the addition of the drug cocktails. TDP43-A315T mice fed NutraGel^®^ lived substantially longer than the mice on MediGel^®^, but we continued to observe GI abnormalities.

Although it is clear that the GI abnormalities in the TDP43-A315T mice we used are a consequence of mutant gene expression in neurons of the gut [17], we do not know whether the mechanism of neuronal dysfunction in these neurons is related to ALS-associated disease processes in cortical or spinal motor neurons. None of the drug combinations we tested were effective in mitigating the GI issues and all treated mice continued to present with both GI abnormalities with hindlimb paresis or paralysis. The degree to which potentially painful GI abnormalities contributed to the appearance of hindlimb weakness is unclear. We did identify a few TDP43 mice fed NutraGel^®^ diets that had clear intestines with hindlimb paresis and paralysis. Notably, in these paralyzed TDP43 mice we did not detect obvious motor neuron loss, which is a defining characteristic of ALS. Thus, even in TDP43-A315T mice exhibiting the expected weakness and paralysis of ALS, the absence of motor neuron loss suggests the model is incomplete.

We had aspired to test many more drug combinations than described here. As we progressed through the experiment and began to recognize the complications of GI abnormalities, we stopped the study after testing 5 combinations. Within these selected combinations we built in some redundancy, building new combinations from a foundation of Riluzole and Edaravone. For example, the addition of Ibudilast to Riluzole and Edaravone had no apparent benefit, and adding additional nutraceuticals to this combination was similarly ineffective. Interestingly, the only combination that showed any promise contained the experimental drug ISRIB (Combo 13). However, it is important to note that this particular cohort had a higher percentage of female mice and thus the modest extension of survival may not be due to drug action since female TDP43 mice tend to live longer. To our knowledge, ISRIB has not been tested in an ALS mouse model. A recent study of two ISRIB-like compounds in G93A SOD1 mice found no benefit [18], while a study of ISRIB in a cell culture model of G93A SOD1 expression in cortical neurons indicated the drug enhanced neuronal survival [19]. Two compounds that act similarly to ISRIB are being pursued for clinical investigation in HEALY ALS. Platform Trial (www.massgeneral.org).

Although we took care to confirm that all of the tested drugs were orally bioavailable, we do not know what level of drug uptake was achieved by delivery in the food. Animals typically feed for several hours after the lights are turned off, with additional short feeding intervals in light [20]. Therefore, the steady-state level of the drug in serum would be much lower than what we measured after oral gavage. Raising the dose of the drug in food could compensate for the more protracted intervals of drug delivery that occur when mixing the drugs with food. However, there may be issues in drug solubility or taste that could be problematic for some compounds. In our experience, delivering the drug cocktails in gel-diets was less stressful for the animals than would occur with frequent gavage.

## Conclusions

The present study examined the utility of the PrP.TDP43-A315T mouse model as a preclinical model for sporadic ALS in trials designed to test combinations of drugs using gel-based diets. Although we observed that TDP43 mice fed gel-based diets lived substantially longer than mice fed standard pellet diets, the gel diets did not completely mitigate gut motility abnormalities that occur in this model. PrP.TDP43-A325T mice on gel diets eventually developed clinical signs of paresis, but the mice lacked evidence of motor neuron loss, which is the hallmark pathology of ALS. Unfortunately, none of the drug combinations we tested acted synergistically to extend survival of the TDP43 mice substantially.

## Acknowledgements

The authors are grateful for the support of Dr. Siobhan Ellison at Neurodegenerative Disease Research Inc., a nonprofit entity. We thank Abhisheak Sharma and Siva Rama Raju Kanumuri of the University of Florida CTSI Translational Drug Development Core for assistance with the pharmacological studies.

